# Mass Photometry of Membrane Proteins

**DOI:** 10.1101/2020.02.28.969287

**Authors:** Anna Olerinyova, Adar Sonn-Segev, Joseph Gault, Cédric Eichmann, Johannes Schimpf, Adrian H. Kopf, Lucas S. P. Rudden, Dzmitry Ashkinadze, Radoslaw Bomba, Jason Greenwald, Matteo T. Degiacomi, J. Antoinette Killian, Thorsten Friedrich, Roland Riek, Weston B. Struwe, Philipp Kukura

**Author notes:** These authors contributed equally to this work (Anna Olerinyova, Adar Sonn-Segev).

## Abstract

Integral membrane proteins (IMPs) are biologically highly significant but challenging to study because they require maintaining a cellular lipid-like environment. Here, we explore the application of mass photometry (MP) to IMPs and membrane mimetic systems at the single particle level. We apply MP to amphipathic vehicles, such as detergents and amphipols, as well as to lipid and native nanodiscs, characterising the particle size, sample purity and heterogeneity. Using methods established for cryogenic electron microscopy, we eliminate detergent background, enabling high-resolution studies of membrane protein structure and interactions. We find evidence that, when extracted from native membranes using native styrene-maleic acid nanodiscs, the potassium channel KcsA is present as a dimer of tetramers – in contrast to results obtained using detergent purification. Finally, using lipid nanodiscs, we show that MP can help distinguish between functional and non-functional nanodisc assemblies, as well as determine the critical factors for lipid nanodisc formation.

## Introduction

Integral membrane proteins (IMPs) constitute 20 to 30% of encoded gene products and have diverse functions, from signalling and transport across membranes to catalysis and mediation of enzymatic reactions^1^. They represent the majority of small molecule drug targets, encompassing G protein-coupled receptors, almost 50% of which are druggable^2^. IMPs are generally more complex to study than soluble proteins because their hydrophobic transmembrane region is intrinsically unstable in aqueous solution, requiring astute strategies to solubilise the protein or maintain a native-like lipid environment. As a consequence, only 1.6% of reported structures in the protein data bank (PDB) correspond to membrane proteins (999 IMP out of 61,301 unique protein structures of <98% sequence identity)^3^.

To overcome the difficulties involved in studying IMP structure and function, a diverse range of membrane mimetic systems are used as protein carriers varying in their usability and capacity to preserve membrane properties. They include detergent micelles, amphipols, nanodiscs and lipid particles^4^. Most often, detergents are used to solubilise and purify IMPs because of their comparative ease of use, making it possible to efficiently extract proteins directly from isolated membranes or intact cells. Detergent micelles, however, do not necessarily mimic the native lipid bilayer, which can affect the function of proteins stabilized in them^5^. Amphipols, whilst more stabilising than detergents, rely on multiple purification steps, increasing the associated experimental complexity. Nanodisc (ND) mimetic assemblies maintain a lipid bilayer *via* annular amphipathic membrane scaffold proteins (MSPs) or styrene-maleic acid (SMA) polymers that encapsulate IMPs in a more native-like environment. SMA lipid particles (SMALPs) and MSP NDs exhibit the improved lateral stability of biological membranes; also, unlike micelles, they do not require excess detergent, and so have a reduced background signal in structural studies^6^.

Membrane mimetic systems (MMS) are inherently heterogeneous, with varying numbers of lipids and other solubilising agents. Coupled with the heterogeneity of IMPs, for example in terms of oligomeric size, studies of IMP structure and function involve challenging and varying combinations of proteins and carriers. This complexity can significantly impact the feasibility, efficacy and ultimate success of IMP studies. As a result, careful purification and characterization of IMP preparations is essential for both functional and structural studies. Current protein purification and characterisation approaches usually rely on a combination of chromatographic techniques; SDS-PAGE; and further analysis by size-exclusion chromatography (SEC), analytical ultracentrifugation, multi-angle light scattering (MALS), negative stain EM or native mass spectrometry (MS) (where applicable). Though well established, these workflows are time-consuming and pose challenges. For instance, MALS characterisation requires a considerable amount of material (>100 ng) and high-resolution separation by SEC. The success of negative stain EM depends to some degree on prior knowledge of likely heterogeneity and sufficient protein size, while native MS of IMPs requires significant expertise and specialised instrumentation. Both of the latter methods necessarily operate under non-native conditions. A method capable of detailed, rapid and accurate sample characterisation in solution could thus dramatically accelerate and improve structural and functional *in vitro* studies of IMPs.

We have recently introduced mass photometry and demonstrated its capabilities in terms of determining molecular mass, resolving different oligomeric states and detecting ligand binding to soluble proteins in a label-free, single molecule sensitive fashion in solution^7^. Given the importance of sample quality, we thus set out to explore the applicability and performance of MP for studying IMPs. Of particular appeal in this context is the universal nature of MP, relying on the detection of changes in the reflectivity of a glass-water interface caused by interference between scattered and reflected light when individual objects bind to the interface^8^ (Figure 1a, **Supplementary Movie 1**). The reflectivity change upon the molecule landing can then be used to determine the object mass using appropriate calibrants (**Supplementary Figure 1**). MP reveals true equilibrium distributions, reports on populations rather than ensembles due to its single-molecule nature, requires minimal sample volumes (µl) and concentrations (<<µM), and does not rely on indirect absorption measurements to determine molecular mass. Moreover, MP naturally extends towards characterising structural heterogeneity^9^ and protein-protein interactions^10^, resulting in a universal platform for studying biomolecules.

**Figure 1.**
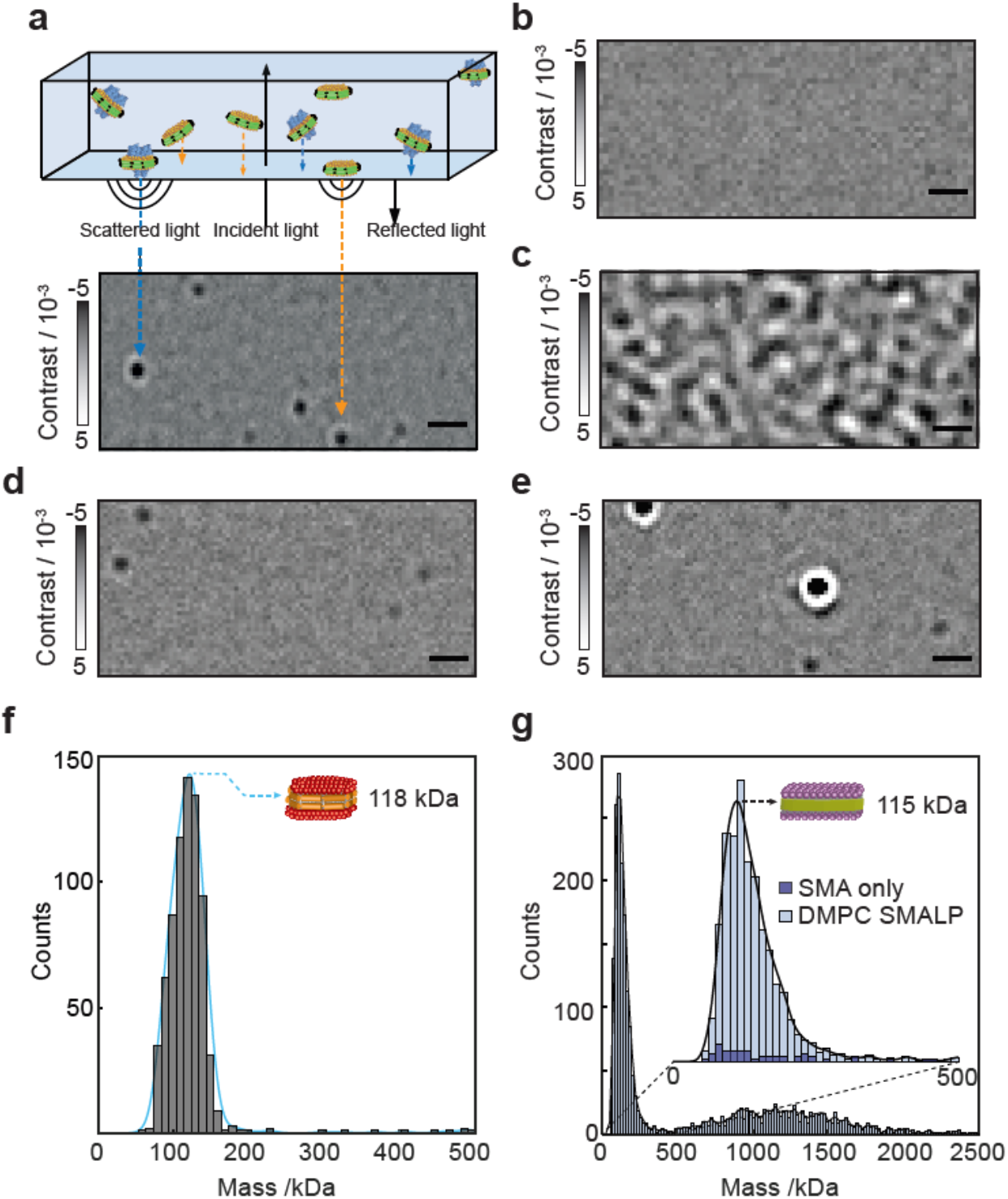
Mass photometry of detergent micelles and nanodisc membrane protein carriers. (**a**) Principle of a mass photometry measurement. Particles land on a glass coverslip, which are detected as a reflectivity change caused by interference between scattered and reflected light. (**b, c**) Representative mass photometry images of imaging buffer and LMNG micelles at 2.5x CMC (0.0025% w/v), respectively. (**d**) Empty MSP1D1 DMPC NDs. (**e**) Empty DMPC SMALPs at a polymer to lipid ratio of 1:1 (w/w). Scale bars: 1 µm. (**f**) The resulting mass photometry histograms and kernel density estimates for empty MSP NDs. (**g**) Corresponding distribution for unoccupied SMALPs (light blue) and SMA polymer aggregates (dark blue).

## Results

### Empty protein carriers

MP is subject to background signatures from nanoscopic empty carriers, because they produce a scattering signal comparable to that of small proteins. To assess the extent to which these could hamper MP measurements, and to help interpret downstream measurements of complex systems, we first studied commonly used membrane mimetic systems in the absence of protein. We began with the detergent lauryl maltose-neopentyl glycol (LMNG) due to its popularity in structural studies based on enhanced stability of membrane proteins solubilised in this manner^11^.

In detergent micelles, using an abundance of detergent above the critical micelle concentration (CMC) aids in solubilising hydrophobic, transmembrane regions of the IMP, thereby maintaining solubility. To assess the effect of an abundance of detergent micelles, we compared LMNG at 2.5x CMC (0.0025% w/v) in buffer (50 mM MOPS, 20 mM NaCl, pH 7.5) compared to a buffer blank (Figure 1b). We observed considerable non-specific detergent interactions (Figure 1c and **Supplementary Movie 2**), due to the effective concentration of LMNG micelles at 2.5x CMC (25 µM)^11^ being an order of magnitude larger than the current optimal concentration range for MP (high pM to mid nM). This caused surface saturation and rapid binding and unbinding on the glass interface, effectively increasing the imaging background, reducing mass resolution and raising the lower detection limit. Other detergents produced similar signatures at concentrations above the CMC (**Supplementary Figure 2**).

**Figure 2.**
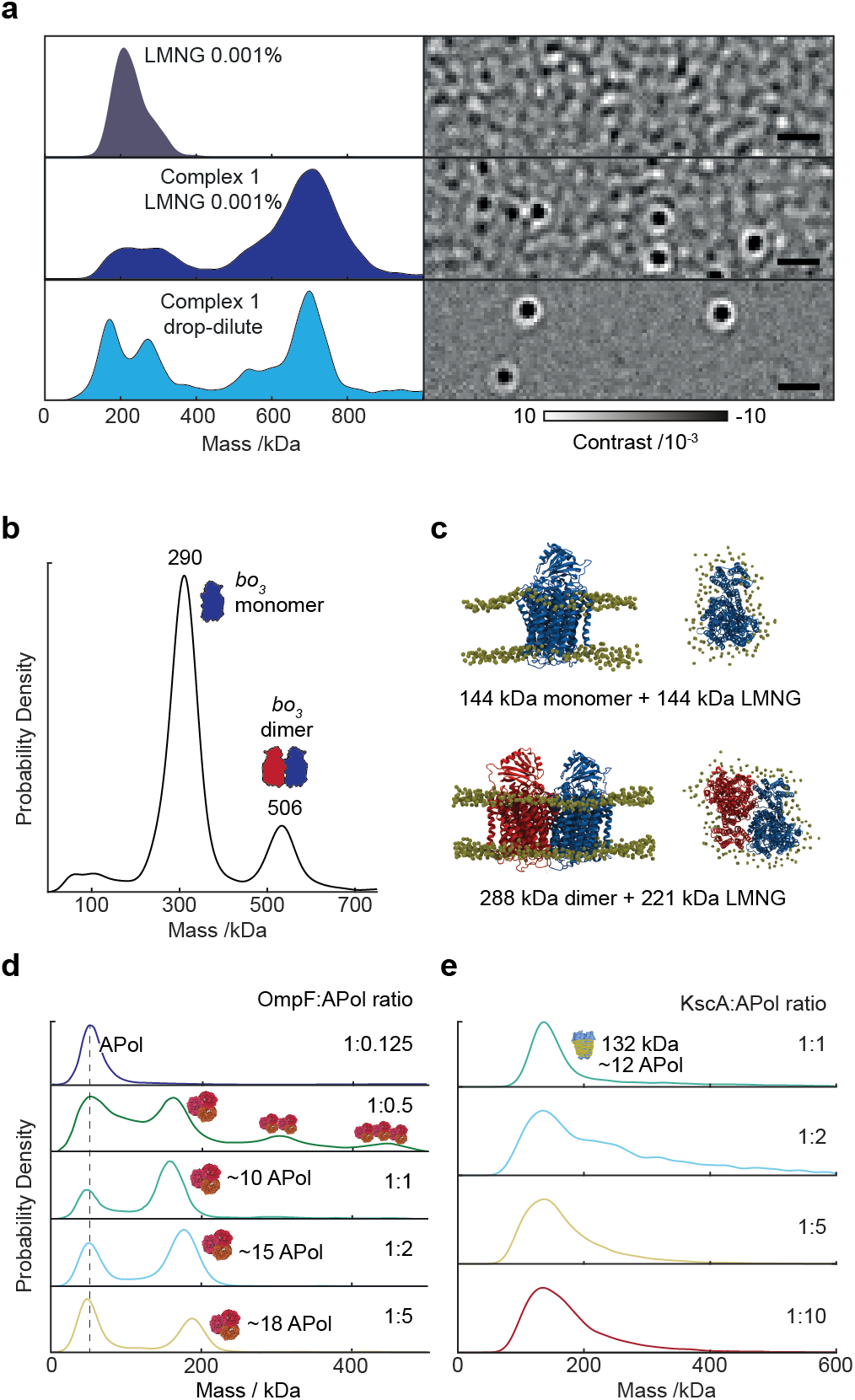
MP analysis of membrane proteins with amphipathic mimetics. (**a**) MP of LMNG at 1x CMC (top) and distributions and typical ratiometric image of respiratory complex I in 1x CMC LMNG (middle) and 2000x LMNG dilution (bottom) after drop dilution. Scale bars: 1 µm. (**b**) Detection of *bo*_3_ oxidase monomers and dimers isolated from LMNG detergent micelles by dilution. Molecular docking and quantitation of bound detergents of *bo*_3_ oxidase monomers and dimers. (**d**) OmpF trimers, and (**e**) KcsA tetramers in amphipols at different protein:APol ratios. KcsA-APol measurements were diluted 200-fold before MP measurement, except for ratio 1:1 which was measured without dilution.

Lipid nanodisc-based systems offer CMC-independent stability, a bilayer-like environment, and potentially lower levels of carrier heterogeneity, as evidenced in our original study^7^. We therefore imaged MSP1D1 DMPC lipid nanodiscs (NDs) and SMALPs (SMA polymer to DMPC lipid ratio 1:1 w/w) using MP. NDs and SMALPs differed from LMNG in that distinct, homogeneous particles were detected (Figure 1d & 1e, **respectively**). Both unoccupied NDs (~118 kDa, Figure 1f) and unoccupied SMALPs (~115 kDa) produced mostly homogeneous mass distributions (Figure 1g). To confirm that we detected unoccupied SMALPs and not disassembled polymer aggregates^12^, we measured SMA polymer at equal concentration as the polymer in unoccupied SMALPs and found a negligible number of particles (Figure 1g, inset, dark blue histogram). Particle distribution analysis revealed that the standard deviation (SD) of the ND peak fit was 23 kDa, or approximately 19%. Compared to standard soluble protein SD of approximately 8-10% of the molecular weight^7^, NDs and membrane mimetics tend to have broader mass distribution due to variations in the number of constituent molecules present. For SMALPs, the SD of the main peak was even larger at 38 kDa, likely arising from varying numbers of lipids and SMA polymers per disc. Unlike MSP NDs, which are limited in size due to the length of the annular belt protein, SMA polymers can in principle form discs of unlimited size, but it is unclear what governs the size of assembled SMALPs^13^. A recent report estimated 140 DMPC lipids per unoccupied SMALP^14^, which would correspond to approximately 20 kDa of SMA polymer and 95 kDa of DMPC lipids on average as detected by MP. In addition to the main SMALP peak, 40% of all observed particles were polydisperse and at higher molecular weight, approximately 500-2000 kDa, indicating some degree of ND oligomerisation. In line with this, it has been reported that SMALPs can form oligomeric ‘rouleaux’ stacks^15^, although this was attributed to a transmission election microscopy (TEM) artefact. This inhomogeneity can also result from an unoptimised SMALP assembly and purification process. MP can therefore guide sample production and purification in SMALP experiments.

### Detergent and APol solubilized IMPs

While background due to unoccupied micelles limits MP performance, we can still distinguish assembled complexes, given sufficient differences in object mass. To illustrate this, we investigated the large *E. coli* protein NADH:ubiquinone oxidoreductase (respiratory complex I, 770 kDa including detergent) in LMNG at 1x CMC (Figure 2a). We could clearly identify assembled protein particles, despite the presence of LMNG, and determine their mass (770 kDa) in excellent agreement with previous studies using analytical ultracentrifugation^16^.

The major remaining limitations were a lack of specificity and detection at low mass in the range of micellar size due to interfering detergent background, as well as a loss of mass resolution. Exploiting the slow off-rate of LMNG molecules from IMPs, gradient-based detergent removal (GRaDeR) has shown great potential to effectively remove empty micelles, while leaving membrane proteins solubilised^17^. We tested a simple 2000-fold drop dilution of complex I from 5x CMC into buffer without LMNG immediately prior to MP measurement. This dilution greatly reduced the detergent background, improving the measurement resolution and enabling us to resolve the low mass peaks. Importantly, the accuracy of the mass measurement was unaffected by LMNG concentration, showing that IMPs can be studied in the presence of LMNG by simply diluting out excess micelles.

We repeated this drop-dilution approach on the *E. coli bo*_*3*_ oxidase (144 kDa monomer), where the presence of *bo*_3_ dimers *in vitro* is debated, with some reports declaring the complete absence of dimers in detergent^18^, and others affirming their existence (although this occurred at the limit of detection, preventing further analysis)^19^. We found monomers and dimers at approximately 80% and 20% relative abundances, respectively, (Figure 2b). Here, the measured monomer mass of 290 kDa suggests approximately 146 LMNG detergent molecules (146 kDa) bound to *bo*_3_, making the simplifying assumption of a similar contrast-to-mass conversion for LMNG to that of protein. Similarly, the measured dimer mass (506 kDa) corresponds to approximately 218 LMNG molecules and a 288 kDa protein contribution. In the aforementioned study affirming the dimers’ existence^19^, *bo*_3_ dimers were suggested to be non-specific artefacts arising from detergent solubilisation. We cannot exclude the possibility that dimer formation is caused by drop dilution from a high LMNG environment, although we did not observe dissociation or significant changes in the relative amount of *bo*_3_ dimers at low concentrations over the course of 360 minutes (**Supplementary Figure 3**). Furthermore, SEC-MALS analysis of *bo*_3_ at higher concentration in 0.003% LMNG (>CMC) was decidedly consistent with our MP analysis – both in terms of molar mass and the relative abundance of monomers and dimers – supporting the occurrence of *bo*_3_ dimers *in vitro* (**Supplementary Figure 4**). Specifically, the conjugate *bo*_3_- LMNG micelle molar mass was 293 kDa (monomer) and 464 kDa (dimer) in 72% and 28% relative abundance, respectively, in good agreement with our results.

**Figure 3.**
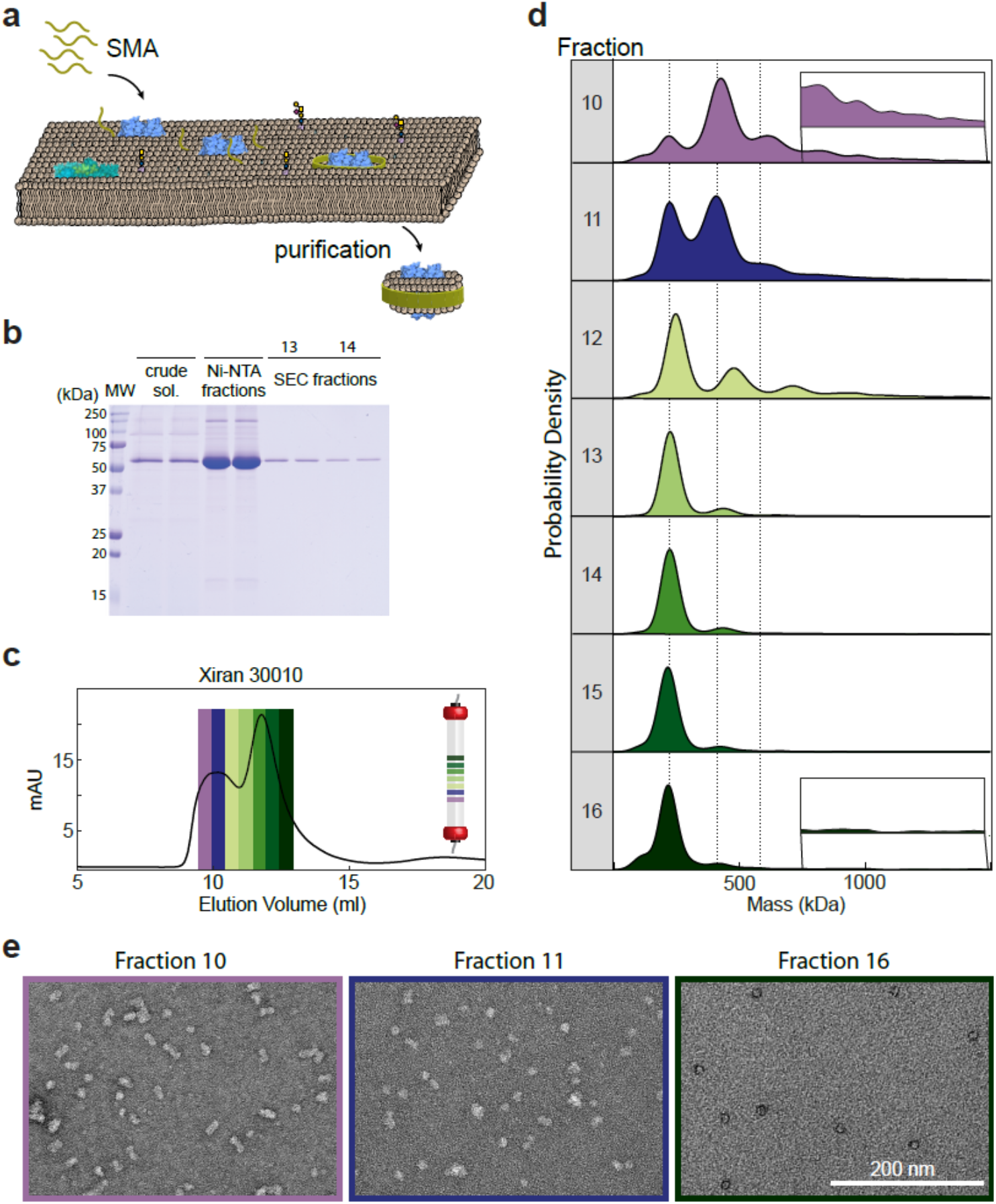
Mass photometry of KcsA in nanodiscs. (**a**) Schematic of native ND formation and extraction of proteins from native membranes by SMA. (**b**) SDS-PAGE of KcsA native NDs (Xiran 30010 SMA) during purification. (**c**) SEC chromatogram of KcsA native NDs, showing the relative protein absorbance at 280 nm. (**d**) MP analysis of SEC fractions 10-16. (**e**) Negative staining EM of fractions 10, 11 and 16 illustrating variability in KcsA native ND assembly.

**Figure 4.**
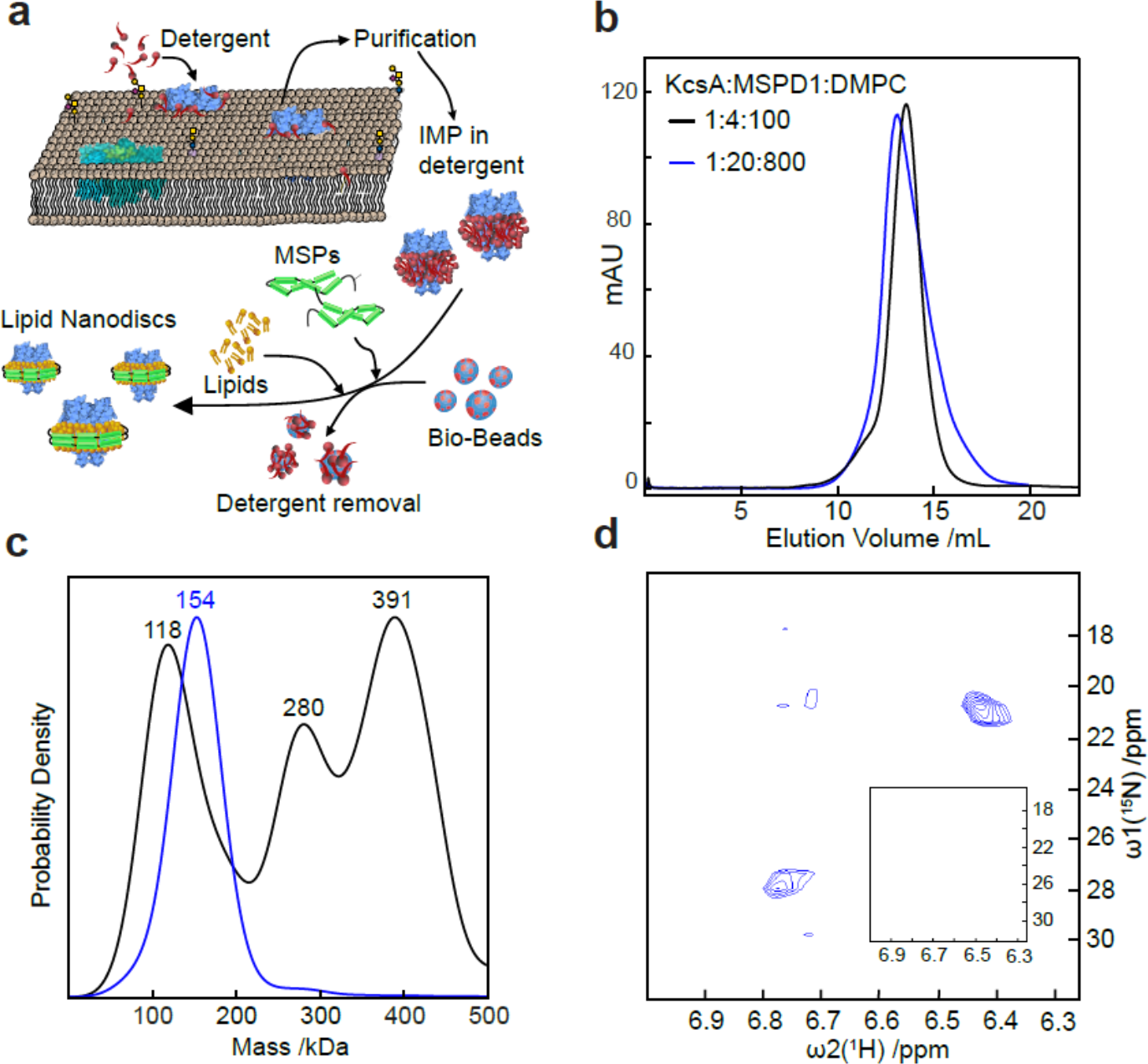
Mass photometry of KcsA in lipid nanodiscs. **(a**) Schematic showing KcsA solubilisation and insertion into lipid nanodiscs. **(b**) SEC chromatograms of KcsA MSP NDs assembled at different KcsA:MSP1D1:DMPC ratios. **(c**) Corresponding MP analysis. **(d**) 2D-HMQC spectra revealing NH4+ binding by KcsA in native NDs prepared in a 1:20:800 KcsA:MSP1D1:DMPC ratio, but no signal was detected from 1:4:100 ratios demonstrating protein inactivity (inset).

To test the feasibility of dimer occurrence, we further explored the *bo*_3_ dimer structure using protein-protein docking in conjunction with molecular dynamics^20^. We simulated the monomer (PDB: 1FFT)^21^ in a POPC bilayer for 60 ns, and used this equilibrated structure to generate a pool of possible dimeric arrangements (best candidate is shown in Figure 2c). We then calculated the transmembrane solvent accessible surface area (SASA) of monomeric *bo*_3_ (21800 Å^2^). Given the mass of LMNG determined by MP (146 kDa), the ratio of SASA per kDa of detergent is 151.4 Å^2^ kDa^−1^, which would amount to the presence of 144 lipids. Assuming this ratio is constant between monomeric and dimeric *bo*_3_, we derived the mass of LMNG bound to the dimer using, as a consensus value, the mean of dimeric model transmembrane SASA (33500 Å^2^). We obtained an LMNG mass of 221 ± 3 kDa, a prediction remarkably close to the experimental value of 218 kDa. These data support the presence of the *bo*_3_ oxidase dimer *in vitro* (Figure 2c), and illustrate the accuracy of MP in quantifying not only the mass of the polypeptide, but also the detergent mass of solubilised membrane proteins.

Amphipols (APols) represent an alternative to detergents for amphipathic IMP solubilisation. APols are small polymers (~4.3 kDa) that function similarly to detergents, adsorbing at hydrophobic patches on IMPs. APols, however, are more stabilising than detergents and excess APol is not necessarily required to solubilise IMPs, meaning background signals from free APol can be minimised for structural and biophysical methods^22^. Nevertheless, their use requires an initial detergent solubilisation step, adding experimental complexity compared to detergent solubilisation alone. MP observations of APols alone were similar to observations of detergent micelles (**Supplementary Movie 6**).

To examine the applicability of MP to APols, we chose the *E. coli* outer membrane protein F (OmpF), which has previously been studied by small-angle neutron scattering (SANS) in APols^23^. OmpF assembles into trimers of β-barrel monomers and functions as a small molecule pore with an overall mass of 111 kDa (**Supplementary Figure 5**)^24^. Mass photometry revealed ratio-dependent assembly of OmpF trimers solubilised in A8-35 APols from 1:0.125 to 1:5 OmpF to APol ratios (Figure 2d). At or above 1:1 OmpF to APol, MP yielded single peaks of OmpF trimers, suggesting predominantly homogeneous samples. As we increased the APol concentration, we observed a clear mass increase corresponding to larger numbers (approximately 10-18) of bound APols. At sub-stoichiometric ratios, however, we found evidence for dimers and trimers of OmpF oligomers (1:0.5), consistent with published SANS results^23^. These results suggest an early onset of aggregation correlated with loss of APol from OmpF-APol trimers, leading to nonspecific hydrophobic protein-protein interactions of OmpF trimers. At the extreme 1:0.125 (OmpF:APol) ratio, we could no longer detect any solubilised protein, likely due to large-scale aggregation. Importantly, loss of OmpF during exchange from detergent to amphipols was minor: from 1:0.5 to 1:5 protein:APol ratio based on UV-Vis measurements (**Supplementary Figure 6**). By contrast, a similar titration for the *Streptomyces lividans* potassium channel, KcsA, did not reveal signs of further oligomerisation (Figure 2e). Instead, we observed loss of KcsA during transfer from detergent to APol below 1:2 protein to APol ratios, as observed by UV-Vis measurements, which can be rationalised by partial disassembly for KcsA tetramers and protein aggregation (**Supplementary Figure 7**).

### Native Nanodiscs

SMA polymers have attracted considerable interest because they can spontaneously solubilise lipid membranes without the need for detergent^25^ (Figure 3a). In short, SMA polymers solubilise IMPs directly from their native environment, retaining natural lipids that may be important for protein stability, assembly, function and interactions. The heterogeneity inherent in native membranes, *i.e.* lipid populations, dynamics., oligomerisation and proximity of different proteins, may be better reflected in extracted native NDs. This is opposed to MSP NDs, where IMPs are solubilised in detergent prior to assembly with known lipids and protein belts, and where their limited diameter may restrict applicability to larger complexes.

To explore MP’s potential for characterising native ND preparations and aiding in structural and functional studies, we expressed *E. coli* KcsA tetramers (80 kDa) and solubilised the membrane using SMA (Xiran 30010). KcsA native NDs were isolated by affinity chromatography and further purified by SEC. Analysis by SDS-PAGE gel (Figure 3b) indicated relatively pure protein preparations, in particular in SEC fractions 13-14. The SEC profile exhibited two main peaks (Figure 3c) and MP analysis of individual fractions taken across both distributions revealed a significant variability in sample heterogeneity (Figure 3d, **Supplementary Figure 8**). Fraction 10 revealed a main peak of ~400 kDa, with additional peaks at ~250 and 600 kDa, indicating a variation in sample size, which may in part be a result of clustering of native nanodiscs or extraction of higher KcsA oligomers. Successive fractions showed a progressive decrease in the number of species, with the most homogeneous fraction (14) eluting at the apex of the second major SEC peak. Fractions 15 and 16, however, were more heterogeneous with a smaller mass shoulder (~125 kDa) next to the main peak (~250 kDa), consistent with minor quantities of empty SMALPs.

To better understand the oligomerisation and composition of the KcsA native NDs, we analysed our preparation’s lipid-to-protein ratio. After affinity purification of the His-tagged KcsA in native NDs, we found that ~44 lipids were present per KcsA tetramer (**Supplementary Figure 9**). Considering a particle size of ~250 kDa, and assuming a polymer contribution of ~30 kDa^26^, the remaining lipid-to-protein ratio would be ~200 if the nanodiscs consisted of a single tetramer per particle, but ~42 if the nanodiscs were composed of clusters of protein, *i.e.* dimers of tetramers (**Supplementary Table 1**). Though the SEC-purified fractions contained only a limited amount of material, we were able to estimate a lipid-protein ratio in the range of ~20-50 from fractions 13 and 14, consistent with KcsA being present as dimers of tetramers, in agreement with signatures of clustering reported previously^27–29^. Additionally, we qualitatively confirmed the observed transition from heterogeneous to homogeneous sample composition by measuring fractions 10, 11 and 16 with negative stain electron microscopy (Figure 3e), which showed larger species in fractions 10 and 11, and a homogeneous distribution of smaller species in fraction 16.

### Lipid Nanodiscs

The high resolution of MP also enabled us to screen and optimise the conditions for reconstitution of KcsA from *S. lividans* into MSP NDs. After selecting the lipid entity and the MSP variant, there are three main variables in the preparation ofKcsA embedded nanodiscs: the absolute and relative concentrations of KcsA, MSP1D1 and the lipids (Figure 4a). In the initial screen (condition B in **Supplementary Table 2**), a KcsA:MSP1D1:DMPC lipid ratio of 1:4:100 was selected on the basis of an excellent size exclusion profile (Figure 4bf, **black**). The MP profile clearly indicated three different species with masses of 118 kDa, 280 kDa, and 391 kDa corresponding, respectively, to empty ND, ND with two KcsA tetramers and two fused NDs (Figure 4c, **black**). By contrast, a KcsA:MSP1D1:DMPC lipid ratio of 1:20:800 exhibited a slightly poorer size exclusion profile (Figure 4b, **blue**), but a much more homogeneous MP profile (Figure 4c, **blue**), with a mass of 154 kDa corresponding to a single KcsA tetramer embedded in a nanodisc consisting of 2 MSP1D1 proteins and approximately 25 lipids. The latter preparation exhibited ^15^NH_4_^+^ ion binding upon addition of 50 mM ^15^NH_4_^+^, as revealed by the presence of 2 strong and 2 very weak NH_4_^+^ cross peaks in a 2D [^15^N,^1^H]-HMQC NMR spectrum assigned to the 4 ion binding sites in the selectivity filter of KcsA (Figure 4d). By contrast, we could not detect any signatures of ion binding in the former sample preparation shown to be highly heterogeneous and lacking KcsA tetramers by MP, demonstrating the lack of functional KcsA (Figure 4d, **inset**).

To further evaluate which of the possible assembly ratios is essential for ensuring assembly and purification of KcsA tetramers, we carried out a screen to determine the optimal IMP to scaffold (mol/mol) ratio with a fixed scaffold:lipid ratio of 1:40 at a lipid concentration of 16 mM. We monitored sample absorbance at 280 nm, indicative of KcsA content, for the following conditions: the initial assembly mixture, the assembly mixture after overnight incubation, the washing fraction from batch Ni^2+^-NTA chromatography, and the elution fraction of purified KcsA nanodiscs (**Supplementary Figure 10**). Characterising all resulting KcsA MSP ND samples by mass photometry revealed homogeneous samples with a consistent mass around 154 kDa (**Supplementary Figure 11**). These results suggest that the IMP:scaffold molar ratio is not crucial for correct sample assembly. Instead, the key parameter controlling the functionality of KcsA embedded in MSP NDs is the scaffold:lipid ratio, contrary to previously published protocols^30^, highlighting the potential of MP in future nanodisc studies.

## Discussion

These results demonstrate that MP can accurately characterise membrane protein carriers and IMPs at the single particle level in solution. Given its universal applicability, speed, and ease of operation, MP is likely to have a significant impact on *in vitro* studies of IMPs. While many of the results from both empty and filled amphipathic carriers can be obtained with alternative techniques at the ensemble level, none can match the speed (60 s per experiment), low sample requirements (<pmoles), and single particle sensitivity and resolution of MP. For SMALPs, we showed the dramatic difference in sample information as revealed by SEC compared to MP. Such sub-chromatographic resolution characterisation of samples during purification holds great promise to improve and accelerate sample preparation for subsequent structural or functional studies. In the case chosen (Figure 3), for example, fraction 15 would be immediately suitable for subsequent characterisation by cryoEM, providing a route to experimentally test whether detergent and native environments are indeed indistinguishable^31^, although the fact that our SMALPs consist almost exclusively of dimers of tetramers already points towards non-negligible differences between the two. The stark differences between the information content from SEC and MP are further illustrated by our results on KcsA in MSP NDs. Two almost indistinguishable SEC profiles were shown to consist of completely different assemblies, with only one, the one identified by MP, being functional.

Despite MP’s advantages, limitations remain: (1) MP’s concentration range is limited to the <100 nM regime, which represents a challenge for the majority of detergents. This can be addressed by borrowing from already proven approaches for translating single molecule techniques from the nM to the µM or even mM regime^32^, or surface passivation approaches. (2) The output of MP is not protein-specific. Gaining insight on protein vs lipid mass thus requires some *a priori* knowledge of the protein mass. (3) The masses reported herein are derived from a calibration based on globular polypeptides because well-defined mass standards – consisting of lipids only, or mixtures of lipids and polypeptides – do not exist, to our knowledge. As a result, we have no way to strictly confirm the accuracy of our mass measurements. Nevertheless, we have previously found very close correlation between MP-measured and predicted mass changes as a consequence of lipid changes alone^7^, and so we expect our results here are also accurate. In future, a more quantitative scale could be provided by obtaining robust standards, or by explicit comparison with native mass spectrometry measurements, assuming, where possible, that no lipids or weakly associated species are lost during the measurement.

Our results should not, however, be viewed exclusively in the context of comparison with existing sample characterisation techniques. Existing techniques are very powerful, and have enabled and continue to yield detailed information on sample composition and heterogeneity. Key to MP is the combination of speed and simplicity, as well its broader capabilities, which include detection of small ligand binding^7^, quantitative evaluation of binding affinities and kinetics in a surface free manner^10,33^, nucleic acid interactions^34^ and more general capabilities for characterising sample heterogeneity^9^. All of these measurements are performed in the same way as the ones presented herein: by adding small volumes (µl) of low concentration (<µM) unlabelled samples in a buffer of choice to a microscope cover slide. This combination of speed and ease of use with a broad feature palette of additional capabilities will make MP a powerful, universal method to study membrane protein structure, function and interactions.

## Methods

Methods and associated references are available in the online version of the paper.

## Supporting information

Movie of empty nanodiscs measured by MP

Movie of 0.0025% LMNG measured by MP

Movie of 3mM DPC measured by MP

Movie of 0.02% DDM measured by MP

Movie of 1% OG measured by MP

Movie of amphipol measured by MP

## Supplementary Information

Supplementary Figures 1-11, Supplementary Tables 1-4 and Supplementary Movies 1-6.

## Acknowledgment

P.K. and A.S.S. are supported by an ERC Consolidator Grant (PHOTOMASS 819593). A.O. is supported by an EPSRC DTC Studentship. Jo. G. is supported by a Junior Research Fellowship at the Queen’s College, Oxford. M.T.D is supported the EPSRC fellowship EP/P016499/1. W.B.S. was supported by Refeyn Ltd. The work in the T.F. laboratory was supported by the Deutsche Forschungsgemeinschaft (DFG) grant 278002225/RTG 2202. We thank Catherine Lichten for feedback.

## Author contributions

Concept: W.B.S., P.K. Methodology: A.O., A.S.S., Jo. G., W.B.S., P.K. Investigation: A.O., A.S.S., Ja. G., D.A., R.B., C.E., J.S., A.H.K., L.S.P.R. and M.T.D. Formal Analysis: A.O. and A.S.S. Writing – Original Draft: W.B.S. and P.K. Writing – Review & Editing: All authors. Visualisation: A.O., A.S.S., W.B.S. and P.K. Supervision: Jo. G., J.A.K., T.F., A.S.S., R.R., W.B.S. and P.K.

## Methods

### 1. Mass Photometry

Mass photometry data was acquired in microscope flow chambers. All microscope coverslips (No. 1.5, 24×50 and 24×24 mm^2^, VMR) were cleaned by sonication with 50% isopropanol (HPLC grade)/Milli-Q H_2_O, followed by sonication in Milli-Q H_2_O (5 minutes each). Microscope coverslips were dried by either a clean nitrogen stream or in the oven at 110 ^°^C for an hour. A small proportion of coverslips was cleaned by applying a layer of First Contact Polymer Optics Cleaner onto the surface, letting it dry for 15 minutes, peeling off the solidified layer, followed by rinsing in ethanol (HPLC grade)/Milli-Q H_2_O and drying with a clean nitrogen stream. Flow chambers were assembled from the clean coverslip immediately after the cleaning process, using double-sided-sticky tape (3M)^7^, and stored prior to use for up to three weeks. MP measurements were performed at a range of concentrations ~10-50 nM, with the exact concentration specified in **Supplementary Table 3**. Where the protein stock concentration was higher, diluting to nM was done immediately prior to the measurement (unless stated otherwise).

For each MP measurement, a buffer solution was added to the flow chamber and the focus position identified and secured for the entire measurement using a focus feedback loop based on total internal reflection of a reference laser beam^8^. Each measurement was taken for either 60 or 90 seconds after ~15 µL of the diluted sample was introduced into the flow-chamber.

All measurements were performed on three similar mass photometers. Most of the data was acquired using a home-built mass photometer as described previously^7^. Briefly, the output of a 520 nm laser diode (Lasertack) was collimated and sent through a pair of acousto-optic deflectors (AODs, AA Optoelectronic DTSXY-400). A 4f telecentric lens system images the deflection by the AODs into the back focal plane of the microscope objective (Olympus UApo N, 100x, 1.49 NA). The objective collects light reflected at the interface between a glass coverslip and some of the light scattered by the sample, with efficient separation of illumination and detection achieved through the combination of a polarising beamsplitter and quarter-wave plate (Thorlabs). The same telecentric lens system images the back focal plane of the objective onto a partial reflector made from a thin layer of silver of 2.5 mm diameter deposited onto a window, which selectively attenuates the reflected light compared to light from point scatterers at the surface by a factor of about 1000. A final lens images the sample onto a CMOS camera (Ximea, MC023MG-SY) with 277.8x magnification, resulting in a final pixel size of 21.1 nm/pixel. Before saving each movie file, areas of 4×4 pixels were binned for an effective pixel size of 84.4 nm/pixel, and frames were averaged 5-fold in time. The entire setup was constructed on a thick (50 mm) metal aluminium plate, and fully enclosed to minimise the influence of air currents.

The *bo*_*3*_ oxidase and respiratory complex I data was acquired with a similar, but commercial mass photometer, One^MP^ (Refeyn LTD, Oxford, UK), with effective pixel size and frame rate as described above. Empty SMALP and OmpF-APol data were acquired on a home-built mass photometer which uses 445 nm Laser diode. Here, the pixel size is 23.4 nm and frame rate is 1 kHz. Prior to saving the images, a 5fold time average and 3×3 pixel binning were applied, resulting in effective frame rate of 200 Hz and effective pixel size of 70.2 nm. Data acquisition was performed using either custom software written in Labview (for the home-built mass photometers) or AcquireMP (Refeyn LTD, v1.2.1) for the commercial instrument.

#### 1.1 Image Processing

##### Mass photometry landing assays

The videos of proteins binding to the glass surface were analysed with DiscoverMP (Refeyn Ltd). The software detects binding events and determines the respective interferometric scattering contrasts. The user can choose how many frames are averaged for continuous background removal (*n*_*avg*_) and can set the thresholds *T*_*1*_ and *T*_*2*_ for the two image filters, which are used to detect the binding events.

Filter 1 is based on T-tests of the pixel intensity fluctuations. As a particle (i.e. IMP) binds to the glass, the pixel intensity changes suddenly. This change is associated with an increase of the filter 1 score calculated as –ln(*p*), where *p* is the *p*-value of the T-test comparing pixel values at *n*_*avg*_ frames before and *n*_*avg*_ frames after the event. The smallest intensity jump amplitude that exceeds random noise fluctuations and is associated with a binding event is controlled by the value of threshold *T*_*1*_.

The signatures of the binding events in interferometric images are radially symmetric. Filter 2 measures the radial symmetry of all pixel neighbourhoods of the interferometric images^35^. The lowest symmetry score expected at the centre of a peak is defined as threshold *T*_*2*_.

Pixel clusters that exceed both thresholds *T*_*1*_ and *T*_*2*_ are used for peak fitting. The amplitude of the peak fit provides an estimate for the interferometric peak contrast. The peak signature (point spread function) is modelled as a superposition of two Sombrero functions multiplied by two Gaussians^7^.

#### 1.2 Calibration Procedure

Contrast-to-mass (C2M) calibration protocol included measurement of several different protein oligomer solutions, with known masses. Each MP calibration was analysed using DiscoverMP, where mean contrast values of all peaks were determined in the software using Gaussian fitting. The mean contrast values were then plotted against the known mass of the proteins (**Supplementary Figure 1** and fitted to a line, *y* = *bx*, with *y* – contrast, *x* – mass and *b* – C2M calibration factor. For the data shown in Figure 4b-c, and **Supplementary Figure 11**, calibration was performed on the basis of mass measurements of empty nanodiscs (118 kDa) in Figure 1f, and a separate measurement of a functional KcsA ND preparation (154 kDa).

### 2. Integral Membrane Protein Preparation

#### 2.1 MSP Nanodiscs

##### Potassium channel KcsA

KcsA from *Streptomyces lividans* was produced using a pET-28a vector containing the coding sequence of the wild-type protein fused to a thrombin-cleavable N-terminal His-tag (monomer 19.9 kDa)^36^. KcsA was expressed in the *E. coli* strain BL21 Star (DE3) (Invitrogen) in Luria-Bertani media. Cells were grown at 37 °C until OD_600_=0.8 and protein production was initiated by addition of 0.5 mM IPTG (Invitrogen). The culture was incubated for additional 5 hours at 37 °C. After protein expression, the cells were lysed by two microfluidizer (Microfluidics) cycles. KcsA was extracted from the membrane with 20 mM dodecylphosphocholine (DPC) detergent by gentle stirring at 4 °C overnight. The cleared extraction mixture was loaded on a Ni^2+^ Sepharose 6 Fast Flow resin (GE Healthcare). After washing the resin with 3.3 mM DPC, KcsA was eluted with 300 mM imidazole and 3.3 mM DPC. Fractions containing the protein were pooled together and the buffer was exchanged to 20 mM Tris-HCl pH 7.4 and 3 mM DPC using a PD-10 (GE Healthcare) desalting column.

##### Expression of membrane scaffold protein (MSP1D1)

MSP1D1 fused to a TEV (Tobacco Etch Virus) protease cleavable N-terminal His-tag was encoded in a pET-28a vector. The MSP1D1 variant of MSP1 deletes the first 11 amino acids of the original MSP1 sequence^37^. Expression and purification of the protein was carried out as described previously^38^. The His-tag on MSP1D1 was cleaved by addition of TEV protease for 16 hours at room temperature. MSP1D1 without His-tag has a molecular weight of 22.0 kDa.

##### Reconstitution of KcsA into DMPC Nanodiscs

**1**) Following the protocol of Shenkarev et al.^39^ KcsA was incorporated into saturated 1,2-dimyristoyl-sn-glycero-3-phosphocholine (DMPC) (14:0) MSP1D1 nanodiscs. DMPC lipids were first solubilised in sodium cholate (cholate/DMPC 2:1 molar ratio). The purified KcsA sample was mixed with MSP1D1, DMPC, and sodium cholate at a molar ratio of 1:20:800:1600 in a buffer containing 20 mM Tris-HCl pH 7.4 and 150 mM NaCl. The final concentration of KcsA was 15 μM. The mixture was incubated overnight at 27 °C while shaking at 150 rpm. Incorporation of KcsA into nanodiscs was initiated by addition of 80% w/v Bio-Beads for 2 hours at 27 °C while shaking at 150 rpm. Empty nanodiscs were separated from nanodiscs containing N-terminal His-tagged KcsA by Ni^2+^ affinity chromatography. The fraction of nanodiscs containing KcsA was eluted with 300 mM imidazole and used for mass photometry measurements. **2**) DMPC lipids were purchased from Avanti and solubilised in 200 mM sodium cholate at a concentration of 100 mM. Lipid nanodisc assembly mixtures containing membrane protein KcsA (tetramer), scaffolding protein MSPdH5 and DMPC lipids were premixed at two different ratios (see **Supplementary Table 2**) to a volume of 500 μL and incubated at room temperature overnight. Lipid nanodisc reconstitution was triggered by the addition of 50% (m/v) Bio-Beads to the assembly mixtures over 4 hours. Assembled nanodiscs were purified with size exclusion chromatography using Superdex 200 chromatographic column (GE Healthcare).

##### Small scale reconstitution of KcsA into lipid nanodiscs

Lipid nanodisc assembly mixtures containing membrane protein KcsA (tetramer), scaffolding protein MSP1D1 and DMPC lipids were premixed at varying ratios to a volume of 50 μL. Concentration of DMPC lipids was kept at 16 mM, concentration of MSP was kept at 400 μM and concentration of KcsA (monomer) was varied in range from 20 μM up to 350 μM. After the overnight incubation at room temperature lipid nanodisc reconstitution was triggered by the addition of 50% (m/v) Bio-Beads to the assembly mixtures over 4 hours. Assembled nanodiscs were purified with batch Ni-NTA chromatography.

#### 2.2 Native Nanodiscs / SMALPs

Commercially available Styrene-Maleic Anhydride (SMAnh) copolymer, Xiran30010 (number-average molecular weight (M_n_) ~2.5 kDa, polydispersity index (PDI) ~2.6), was a kind gift from Polyscope Polysciences (Geleen, NL). Conversion of the SMAnh polymers into the acid form (SMA) was achieved by hydrolysis under base-catalysed conditions as detailed previously^40^. SMA stock solutions were prepared at final concentrations of 5% (w/v).

##### Preparation of E. coli membranes overexpressing KcsA

Total membrane fractions of *E. coli* cells (strain BL21(λDE3)) producing KcsA were obtained as described previously^25^. Briefly, cells were transformed with an N-terminal His-tagged pT7-KcsAvector, containing the *KcsA* gene. Membrane preparations were obtained by differential centrifugation after cell wall lysis and mechanical disruption through a French press. Membrane pellets were resuspended in buffer (Tris-HCl 5 mM, NaCl 300 mM, KCl 15 mM, pH 8) to an OD_600_ of ~4. After lipid extraction according to the method of Bligh and Dyer^41^, the total phosphate content was determined to be 10 mM using the method of Rouser *et al*.^42^. Membrane suspensions were stored at −20 °C until further use.

##### SMA-mediated solubilisation of KcsA from E. coli membranes

Membrane stocks (4.8 mL) were thawed on ice and diluted with solubilisation buffer (Tris-HCl 50 mM, NaCl 300 mM, KCl 15 mM, imidazole 10mM, pH 8) to total volume of 33 mL, resulting in phosphate concentration of ~1.5 mM and protein concentration of ~1 mg/mL. Polymer was added to the suspensions at a final concentration 1% (w/v). The mixture was incubated on a rotary disc at 4°C overnight, during which the suspension cleared up significantly. To remove any unsolubilised material the mixture was centrifuged at ~40,000 g for 1 h at 4 °C to pellet the non-soluble fraction. The solubilised fraction (supernatant) was carefully removed and used for Histag purification of the nanodiscs.

##### Purification of KcsA native nanodiscs

The solubilised fraction containing the nanodiscs was added to 6 mL of HisPure Ni-NTA agarose beads (Thermo Scientific) and incubated on a rotary disc at 4 °C overnight. The beads were then loaded on a gravity-flow column for the affinity purification. First, the flowthrough was collected, followed by washing with increasing amounts of imidazole (10 mM, then 50 mM) and finally elution with a high concentration of imidazole (300 mM). Elution fractions containing pure KcsA were combined and the pooled fractions were concentrated using centrifuge spin filters (Amicon, 15 mL, 10 kDa MWCO). The imidazole was removed by washing the concentrate three times with buffer (Tris-HCl 50 mM, NaCl 300 mM, KCl 15 mM, pH 8).

The isolated KcsA native nanodiscs were further purified and analysed by size exclusion chromatography (SEC) on an ÄKTA pure system (GE Healthcare), using a Superdex 200 10/300 GL size exclusion column (GE Healthcare). Protein was eluted using buffer (Tris-HCl 50 mM, NaCl 150 mM, KCl 15 mM, NaN_3_ 1 mM, pH 8). Fractions were collected on an autosampler and UV detection was performed at λ= 280 nm. Collected fractions were frozen and stored at −80 °C until further use.

##### Determination of sample purity using SDS-PAGE

Samples were incubated with Laemilli buffer without any reducing agent, loaded onto 13% SDS-PAGE gel, and run at 175 V for 1 hour. The gels were stained with coomasie blue and scanned for further analysis.

##### Determination of protein-to-lipid ratios in KcsA native nanodiscs

Protein concentration was determined using Pierce micro BCA protein assay kit (Thermo Scientific). The standard protocol was modified by using 2.5-fold the recommended amount of reagent C, as the SMA copolymers chelate copper and thus an excess is required. Furthermore, 1% SDS was added in order to ensure that the membrane proteins remained in solution. Weight concentration was converted to mole concentration of protein on the basis of a KcsA tetramer of ~80 kDa.

Lipids were extracted according to a modified version of the method of Bligh and Dyer^41^, namely without the use of hydrochloric acid but rather under alkaline conditions to prevent possible aggregation and precipitation of SMA bound to nanodiscs or lipids. The lipid concentration was determined based on total phosphate content according to the method of Rouser *et al*.^42^. For the SEC fractions, two samples had to be combined (fraction 13 and 14) in order to give sufficient material to be above the lower limit of detection.

The contribution of polymers in terms of mass to a single nanodisc is difficult to determine. If we assume two belts of polymers per disc, the total amount of polymer per nanodisc is approximately 30 kDa, or about ten polymer molecules with M_n_ = 3 kDa^26^.

#### 2.3 Detergent Micelles

##### bo_3_ oxidase

*E. coli bo*_3_ oxidase was expressed in *E. coli* strain BL21(DE3) star/pET*cyo* _*his*_*cyoC*. Plasmid pET*cyo* _*his*_*cyoC* was a kind gift from Prof. Christoph von Ballmoos (Bern, Switzerland). Cells were grown at 37 °C in LB medium-supplemented with 0.5% glycerol, 2 mM MgSO_4_, 0.03 mM FeSO_4_ and 0.01 mM CuSO_4_. At an optical density of 1.5, gene expression was induced by an addition of 0.5 mM IPTG and cells were grown for further 2 hours. Cytoplasmic membranes were suspended in buffer A (50 mM MOPS and 20 mM NaCl, pH 7.5) and membrane proteins were solubilised by adding LMNG dropwise to a final concentration of 2% (w/v). Solubilised proteins were loaded onto a Probond Ni^2+^-IDA column (25 mL) equilibrated in buffer A containing 20 mM imidazole and 0.005% LMNG. After washing with 92 mM imidazole, bound proteins were eluted in a single step to 284 mM imidazole. The eluate was concentrated by ultrafiltration (Amicon Ultra-15, 100 kDa molecular weight cut-off) and subjected to size exclusion chromatography on Superose 6 (300 mL) in buffer A containing 0.005% LMNG. The main peak eluting after 177 mL was concentrated to 100 µM, aliquots were shock frozen in liquid nitrogen and stored at 80 °C until further use.

##### Respiratory complex I

Respiratory complex I from *E. coli* was prepared as described previously^43^ with slight modifications. After affinity chromatography on a Probond Ni^2+^-IDA column (35 mL), the complex was subjected to a Superose 6 (24 mL) size exclusion column. Peak fractions were concentrated to 30 µM (Amicon Ultra-15, 100 kDa MWCO) and stored in 50 mM MES/NaOH, 50 mM NaCl, 5 mM MgCl_2_, pH 6.0 with 0.005% LMNG at −80 °C. MP measurements of complex I were performed at 15 nM. Dilution to measured concentrations was performed with buffer A either without any detergents or with 0.001% LMNG.

#### 2.4 Amphipols

##### E. coli outer membrane protein F (OmpF) in APol

OmpF was expressed and purified as described previously^44^ in 20 mM phosphate buffer (pH 6.5, 1% OG) at a concentration of 11 µM. Bio-Beads (Bio-Rad) were used for exchange from detergent to amphipols and detergent adsorption according to Zoonens, *et al*.^22^. Approximately 1 g of Bio-Beads was washed for 25 min with methanol (HPLC grade), 20 min with Milli-Q H_2_O, and 2x 20 min with OmpF phosphate buffer (20 mM phosphate buffer, pH 6.5). Each wash was done at room temperature with gentle agitation. The beads were rinsed twice with Milli-Q in between steps. Fresh A8-35 amphipol (Anatrace) stock solution (10%) was prepared and diluted 10x in OmpF buffer. A range of amphipol to integral membrane protein ratios was calculated, from 0:1 (control) to 5:1, in order to determine the lowest ratio for effective protein extraction. Amphipol solution, integral membrane protein and buffer were added to a final exchange volume of 200 µL. An excess of prepared Bio-Beads was added to each combination, which was then incubated at 4 °C for a total of 16 hours, with a change of BioBeads after 2 hours.

Once detergent removal was complete, the protein – amphipol solution was carefully removed and OmpF – amphipol was buffer-exchanged into OmpF Tris buffer (20 mM Tris, 100 mM NaCl, pH 8.0) using 0.5 mL molecular concentrators with molecular weight cut-off of 30 kDa (Amicon). The sample was spun at 10,000 x g at 4 °C for 2 minutes, after which the flow through was discarded and 300 µL of fresh buffer was added. This was repeated 5 times, checking for protein precipitation after each spin cycle. After OmpF buffer exchange, light absorbance at 280nm was measured for each sample using Nanodrop device in order to determine protein concentration. The samples were subsequently ultra-centrifuged at 150,000 x g for 20 minutes. After centrifugation, absorbance at 280 nm of the supernatant of each sample was measured again using Nanodrop for comparison of the amount of protein retained in the supernatant (**Supplementary Figure 6**). This was used to help determine the minimal amount of amphipol needed to solubilise the membrane protein. Once the insertion was complete, the samples were stored at 4 °C for up to a few weeks.

##### Potassium channel – KcsA in APol

KcsA solubilised in DPC micelles (preparation described above) was exchanged to amphipols using the same protocol described above for OmpF, with a few minor differences; KcsA buffer (20 mM Bis-Tris, 150 mM NaCl, pH 7) was used throughout the whole protocol and a second exchange of Bio-Beads was added after 4 hours of incubation in the amphipol preparation. **Supplementary Figure 7** shows the absorbance values for the different ratio for KcsA-amphipol.

### 3. SEC-MALS

The analysis was performed on an Agilent 1200 HPLC system with an autosampler, diode array detector and differential refractive index detector connected in-line with a 3-angle light scattering detector (Treos, Wyatt Technology). Light scattering data were collected from an injection of 30 μL *bo_3_* oxidase at 87 μM onto a SHODEX KW-803 300 mm x 8 mm column equilibrated in 50 mM MOPS pH 7.0, 300 mM NaCl and 0.003% LMNG at 0.5 mL/min and analysed as a protein conjugate using the Astra V software (v 5.3.4.20). The extinction coefficient of *bo_3_* was predicted from its sequence to be 299 M^−1^cm^−1^ at 280 nm and the detergent assumed to have no absorbance at 280 nm (an extinction coefficient = 0). We measured the refractive index increment, d*n*/d*c*, for LMNG by injecting a series of dilutions of LMNG in water onto our differential RI detector and obtained a value of 0.151 mL/g. In addition, the average d*n*/d*c* for proteins of 0.185 mL/g was used for the analysis.

### 4. Negative Staining Electron Microscopy

For negative staining, grids (300 mesh Cu carbon film) were glow discharged for 20 seconds at 15 mA (Leica EM ACE 200). 10 µL of sample was applied to the grid for 2 minutes, blotted, stained with 2% uranyl acetate for 20 seconds, blotted and allowed to air dry. Images were acquired on a 120kV Tecnai 12 (Thermofisher) TEM equipped with an OneView digital camera (Gatan).

### 5. NMR Spectroscopy

2D [15N, 1H]-HMQC experiments with H_2_O/^15^NH4^+^ defocusing and selective excitation burp pulses were all collected at 305 K on a Bruker 700 MHz Avance III spectrometer equipped with a triple resonance cryoprobe.

### 6. Molecular Dynamics Simulations

We carried out Molecular Dynamics (MD) simulations of *E. coli bo*_*3*_ ubiquin oxidase (PDB: 1FFT)^21^ in a lipid bilayer using GROMACS 2016.4 CUDA^45^. The Amber14sb all-atom force field^46^ was used to describe the protein and the lipid all-atom force field to describe the palmitoyloleoyl-phosphocholine (POPC) lipids. All simulations were run in the NPT ensemble. We first assembled a POPC membrane using packmol^47^ and subsequently equilibrated it in TIP3P water for 20 ns. *bo*_3_ was then inserted into the equilibrated bilayer according to the arrangement predicted by the OPM server^48^. The resulting protein-lipid system was then filled with TIP3P water, charge neutralised with chloride ions and energy minimised *via* steepest descent, with the maximum force tolerance set to 200 kJ mol^−1^ nm^−1^. Finally, the system was simulated for 60 ns, where the first 50 ns were excluded as part of the equilibration, and the last 10 ns used for production.

In all simulations long-range interactions were calculated using the particle mesh Ewald (PME) method and a cut-off of 12 Å was used for van der Waals and Coulombic interactions. The LINCS constraint was used to restrain bonds involving hydrogen atoms. Simulations utilised a 2 fs integration time step, updating the neighbour lists every 10 steps. An atmospheric pressure of 1 bar was maintained *via* anisotropic pressure coupling using a compressibility *k*_x_ = *k*_y_ = *k*_z_ = 4.5 × 10^−5^ bar^−1^, with off-diagonal terms *k*_xy_ = *k*_xz_ = *k*_yz_ = 0 bar^−1^ and time constant τ_p_ = 1.0 ps. The protein, lipids and solvent (water and ions) were individually coupled to a heat bath at 310.15 K with time constant τ_t_ = 0.1 ps.

#### 6.1 Protein-Protein Docking

The prediction of a possible *bo*_3_ dimeric arrangement was carried out using the protein-protein docking software JabberDock 1.0^20^. JabberDock generates protein surface representations (STID maps) simultaneously describing atomic charge, distribution and dynamics, and then produces docking candidates of maximal surface complementarity by exploiting the POW^er^ optimisation engine^49^. To this end, the last 10 ns of MD simulation were extracted to generate a STID map of *bo*_3_ representative of its local dynamics in a lipid bilayer.

The docking starting point featured two *bo*_3_ monomers centred at the origin. One monomer was kept fixed, while the other was allowed to translate in the *xy* plane and to rotate along its *z* axis between 0 and 2π radians. This axis of rotation was also permitted to precess into the *xy* plane between an angle of −0.05 and 0.05 radians. POW^er^ navigated this conformational space, looking for a region associated with two *bo*_3_ monomers of maximal surface complementarity. Optimisation was performed over 200 iterations using 60 randomly initialised particles. The “kick and reseed” procedure involves randomly reinitialising particles that have converged to a local minimum before placing a repulsive potential on the converged site to prevent oversampling. The optimisation process was repeated three times, with memory of the previous landscape including the repulsion potentials kept for future iterations. The docking procedure thus evaluated 36000 docked poses, which were then clustered by *K*-means into 300 representative poses (cluster centroids). The dimer shown in Figure 2c is a representative model of the ensemble, relaxed in a lipid bilayer following the same MD simulation protocol adopted for monomeric *bo*_3_.

The SASA of the equilibrated *bo*_3_monomer and all 300 predicted dimers were calculated using the Shrake-Rupley algorithm as implemented in VMD^50^. The mean and standard deviation of all dimeric models SASA was calculated as consensus value.

### 7. Native Mass Spectrometry

Mass spectra were recorded on prototype Orbitrap Q Exactive UHMR mass spectrometer (Thermo Fisher Scientific), equipped with a Nano Flex nanospray source and offline nanospray source head. OmpF prepared in 100 mM ammonium acetate at 2x CMC octyl β-D-glucopyranoside detergent (Anatrace) was transferred into the mass spectrometer using gold-coated, borosilicate glass capillaries prepared in house. The mass spectrometer was operated in positive ion polarity and in AGC prescan mode with a maximum inject time of 100 ms and target of 1e6. Capillary voltage was 1.4 kV, transfer capillary temperature 80 °C, inject flatapole 10 V, inter-flatapole lens 6 V, bent flatapole 4 V. Ion optics transmission was set to high *m/z*, and detector mode to low *m/z*. Pressure setting was 6-8 V. No insource trapping or in-source activation was applied. Direct HCD voltage applied to remove the detergent micelle was 250 V. Microscans was set to 10, no averaging was applied, and transient time was 64ms, corresponding to a resolution of 12,500 at *m/z* 400. 25 scans were recorded and averaged using the Xcalibur software package v2.2-4.1 (Thermo Fisher Scientific).

