## Supplementary Figures and Tables for "Mass Photometry of Membrane Proteins"

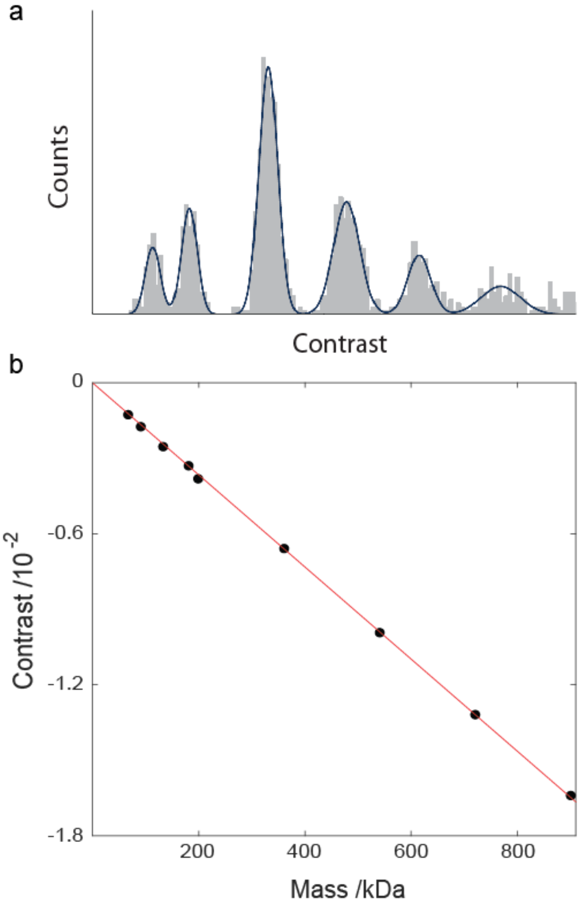


**Supplementary Figure 1. Contrast-to-mass (C2M) calibration. (a)** Example histogram of a mass calibrant as measured by MP. **(b)** The contrast of two proteins of known mass with different oligomeric states were plotted vs their mass (66, 90, 132, 180, 198, 360, 540, 720, 900 kDa). The red line is the linear fit to the data according to $y = bx$, with $b = -1.83 \cdot 10^{-5}$ – C2M calibration factor.

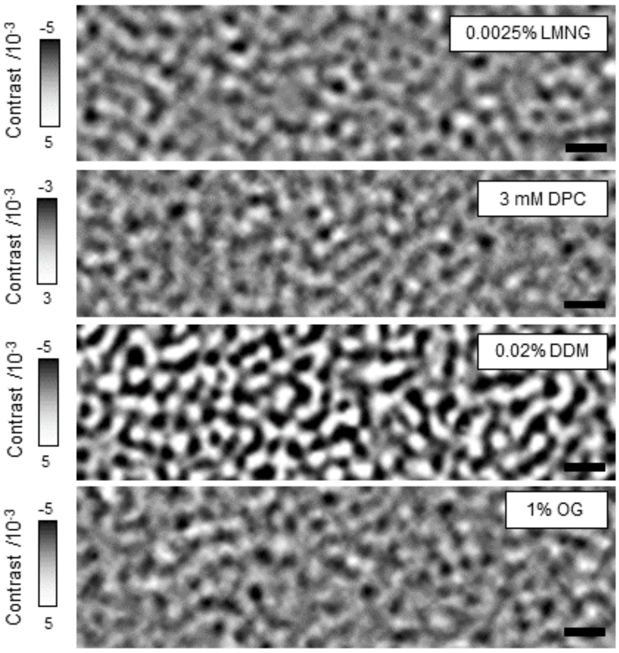


**Supplementary Figure 2. MP of detergent micelles.** Images of detergents LMNG, DPC, DDM and OG at or above the critical micelle concentration (CMC). Accompanying movies are: Supplementary Movie 2 (LMNG), Supplementary Movie 3 (DPC), Supplementary Movie 4 (DDM) and Supplementary Movie 5 (OG). Scale bar: 1 μm.

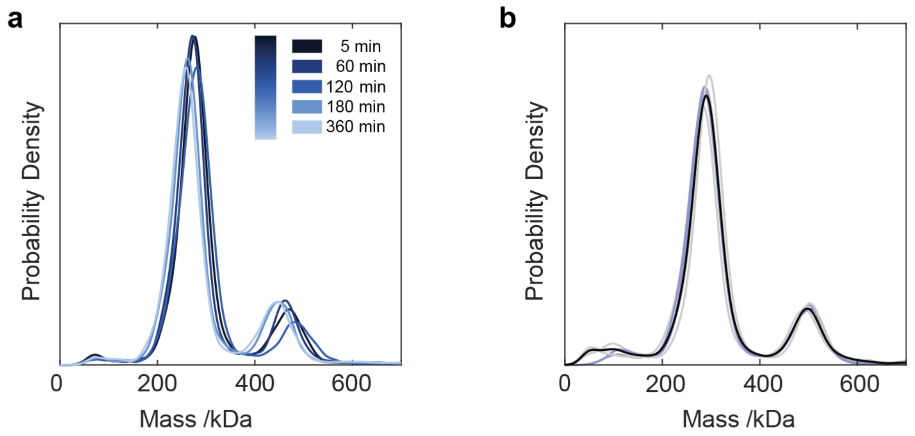


**Supplementary Figure 3: *E. coli bo₃* oxidase reproducibility and stability upon drop dilution.** (**a**) MP measurements of 20 nM *bo₃* oxidase at different time point after drop-diluting in detergent-free buffer. Final LMNG concentration is ~0.001x CMC. Colours correspond to different waiting times after dilution. (**b**) Repeats of MP measurements of 10 nM (light-blue lines) and 20 nM (grey lines) *bo₃* oxidase showing the reproducibility of the measurement. Black line corresponds to the average over all repeats.

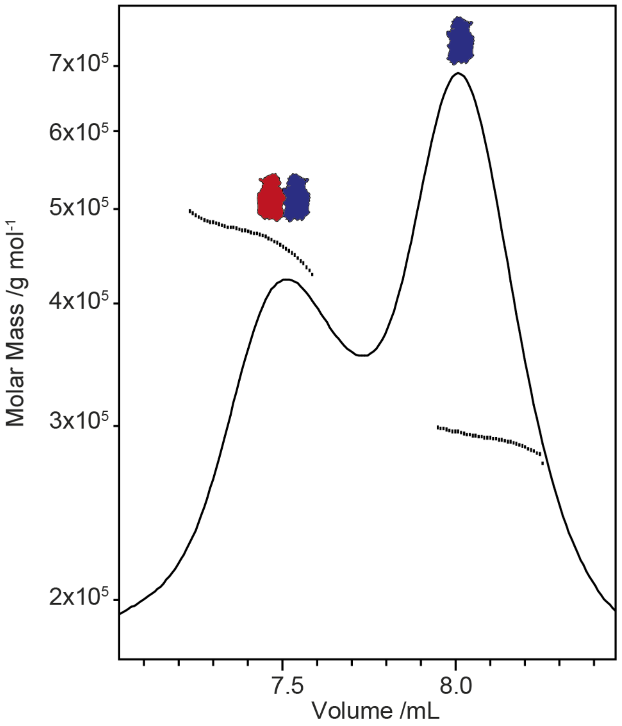


**Supplementary Figure 4: SEC-MALS of *E. coli bo₃* oxidase.** UV absorbance chromatogram and MALS traces (black botted lines) of *bo₃*-LMNG micelle complexes in 0.003% LMNG containing running buffer. The conjugate protein-detergent molar mass was 293 kDa (monomer) and 464 kDa (dimer). The molar mass of individual components were 140 kDa *bo₃* plus 153 kDa LMNG (monomer) and 259 kDa *bo₃* plus 205 kDa LMNG (dimer).

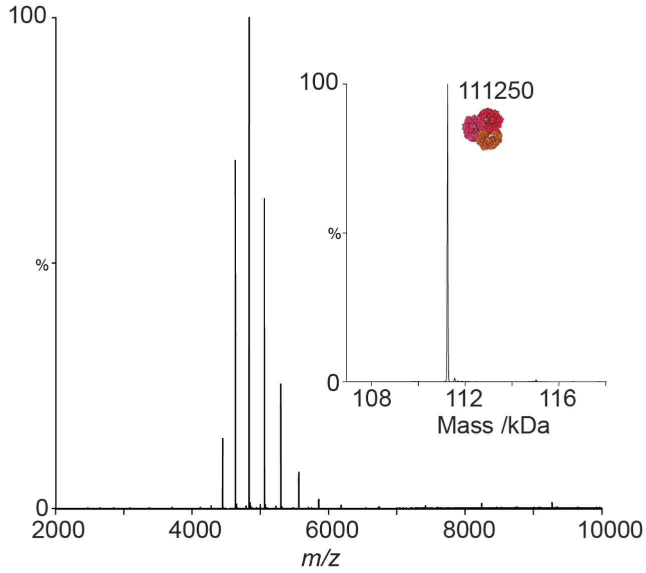


**Supplementary Figure 5. Native mass spectrometry of OmpF**. OmpF at 2 μM measured in 100 mM ammonium acetate with 2 x CMC n-octyl-β-D-glucopyranoside (OG).

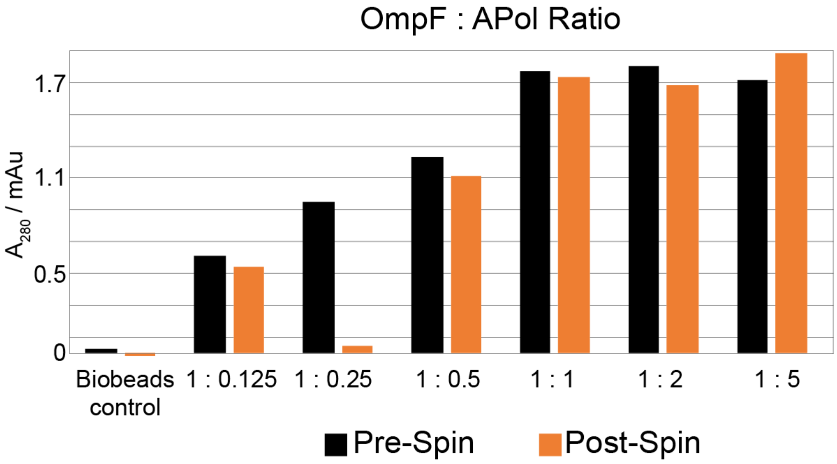


**Supplementary Figure 6. Analysis of OmpF detergent-amphipol exchange**. UV-VIS $A_{280}$ measurement of OmpF prior to ultra-centrifugation step in the amphipol insertion procedure (black) compared to the absorbance of the supernatant post-ultracentrifugation (orange).

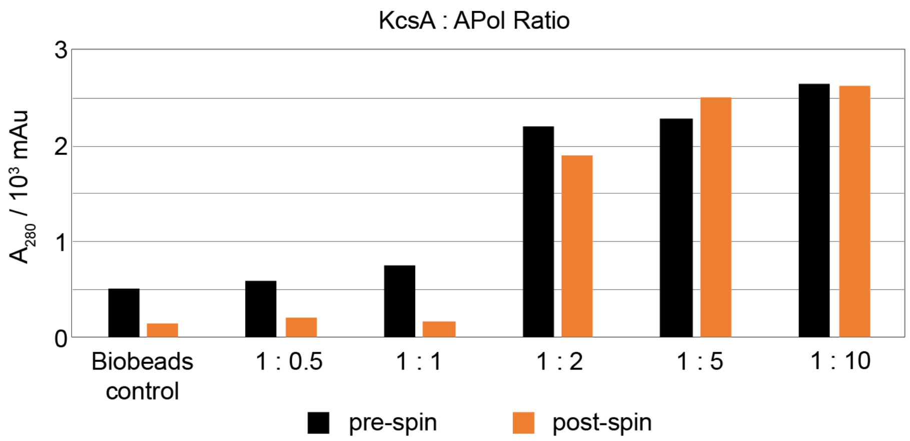


**Supplementary Figure 7. Analysis of KcsA detergent-amphipol exchange**. UV-VIS $A_{280}$ measurement of KcsA prior to ultra-centrifugation step in the amphipol insertion procedure (black) compared to the absorbance of the supernatant post-ultracentrifugation (orange).

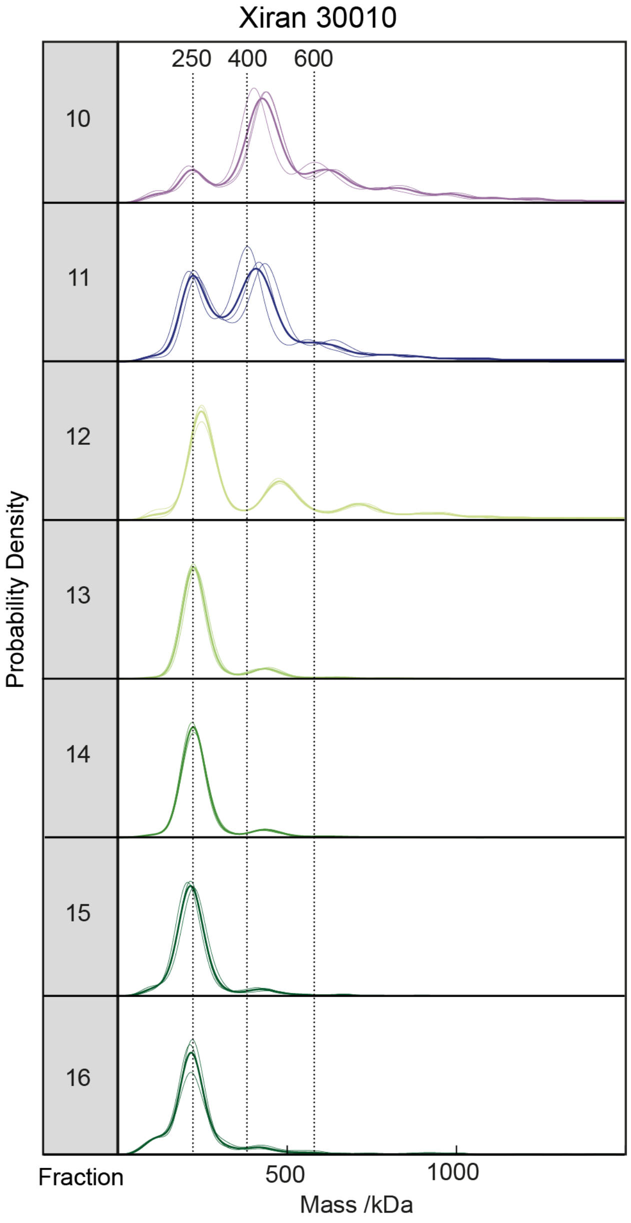


**Supplementary Figure 8. MP replicates of *E. coli* KcsA native ND SEC fractions.** Each fraction was diluted 5-10x and measured immediately three times on separate coverslips. The thick outline for each fraction represents the average data from the three separate experiments.

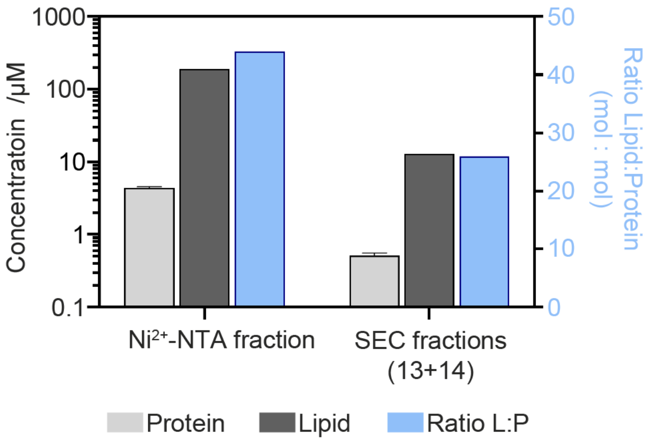


**Supplementary Figure 9. Lipid/protein analysis of *E. coli* KcsA native NDs.** Experimentally derived protein and lipid concentrations and their respective molar ratios for the $Ni^{2+}$-NTA fraction compared to combined fractions 13 and 14 from size-exclusion chromatography.

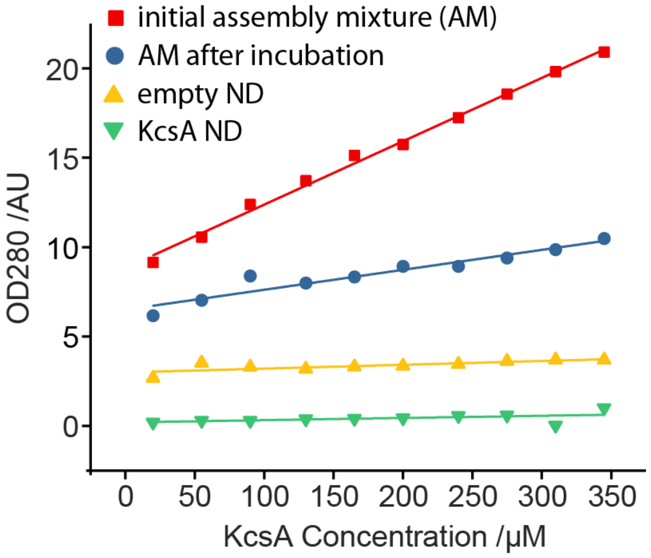


**Supplementary Figure 10. KcsA ND assembly screen via $OD_{280}$.** A series of small-scale assemblies (50 μL each) of potassium channel KcsA into MSP nanodisc was carried out to screen for optimal membrane protein to scaffold ratio. Sample absorbance at 280 nm was monitored for the initial assembly mixture, assembly mixture after overnight incubation, washing fraction from batch $Ni^{2+}$-NTA chromatography and elution fraction or purified KcsA nanodiscs. Recorded absorbances are depicted as a function of the input KcsA concentration.

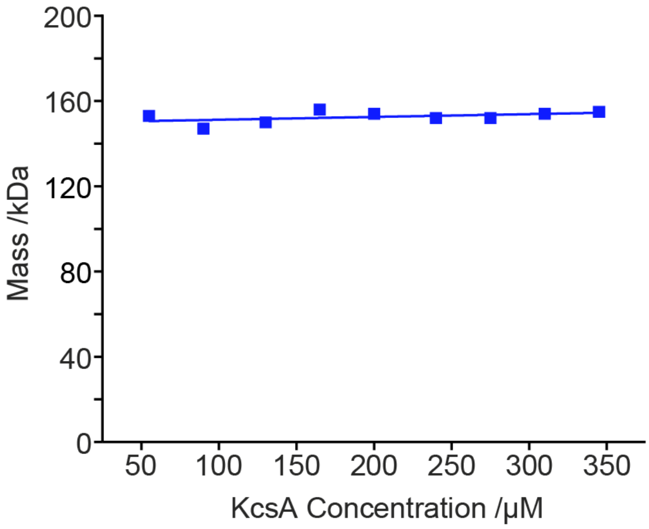


**Supplementary Figure 11. KcsA ND assembly mass distributions.** The average mass of the final nanodisc particle distributions was measured with MP. All of the test assemblies resulted in a homogeneous sample with a consistent mass. This shows that membrane protein to scaffold ratio of up to 1:4.5 does not compromise the sample fitness and therefore is optimal for maximum ND yield per scaffold.

### Supplementary Tables

| ***Theoretical assemblies*** | 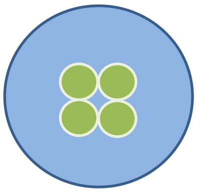 | 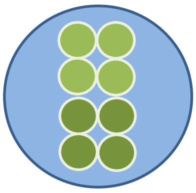 | 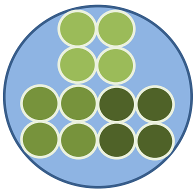 | 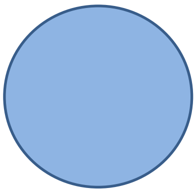 |
|---|---|---|---|---|
| **~Ratio of lipid-to-protein (mol/mol)** | 200 | 42 | 0 | 300 [lipids] |
| **Supramolecular organisation** | Single tetramer | Dimer of tetramers | Trimer of tetramers | Empty SMALP |
| Protein (*) | 80 kDa (1 mol) | 160 kDa (2 mol) | 240 kDa (3 mol) | 0 kDa (0 mol) |
| Lipid (**) | 140 kDa (195 mol) | 60 kDa (84 mol) | 0 kDa (0 mol) | 220 kDa (306 mol) |
| Polymer (***) | 30 kDa (10 mol) | 30 kDa (10 mol) | 30 kDa (10 mol) | 30 kDa (10 mol) |
| Total mass (†) | 250 kDa | 250 kDa | 270 kDa | 250 kDa |

*Protein: KcsA tetramer = 80 kDa. **Lipid: POPE = 0.718 kDa. ***Polymer: $M_n$ = 3 kDa. †Total in one nanodisc.

**Supplementary Table 1: Theoretical lipid/protein/SMA stoichiometry of KcsA native NDs**

| | KcsA tetramer : MSP1D1 : DMPC (molar ratio) | KcsA tetramer / μM | MSP1D1 / μM | DMPC lipids / mM |
|---|---|---|---|---|
| Assembly A | 1:20:800 | 20 | 400 | 16 |
| Assembly B | 1:4:100 | 50 | 200 | 5 |

**Supplementary Table 2: KcsA:MSP1D1:DMPC ND assembly.** Two assembly ratios screened for the reconstitution of the potassium channel KcsA into the lipid nanodiscs

| Protein | Dilution / Concentration |
|---|---|
| KcsA in MSP NDs | 20 nM |
| KcsA in SMALP | 5-10 fold dilution |
| KcsA in APol | ~ 50 nM |
| *bo*$_3$ oxidase | 10 or 20 nM |
| *bo*$_3$ oxidase – time stability assay | 20 nM |
| complex I | 15 nM |
| OmpF in Apol | 30 nM |
| Empty MSP NDs | ~10-50 nM |
| Empty SMALP | 20-40 nM |
| SMA polymer | 2 or 6.7 µg/ml |

**Supplementary Table 3. Protein/carrier concentrations in MP measurements.** Detailed information on the concentration of samples. In cases where the concentrations were not precisely known, a dilution factor was given (i.e. KcsA in SMALPs).

| Protein | Number of binned frames, $n$ | Threshold 1 |
|---|---|---|
| KcsA in MSP NDs | 5 | 1.5 |
| KcsA in native NDs | 5 | 1.5 |
| KcsA in APol | 10 | 1 |
| *bo₃* oxidase | 10 | 0.5 |
| *bo₃* oxidase – time stability assay | 5 | 0.9 |
| complex I | 3 | 1.5 |
| OmpF in Apol | 5 | 1 |
| Empty MSP NDs | 5 | 1.5 |
| Empty SMALP | 3 | 1.5 |
| SMA polymer | 3 | 1.5 |

**Supplementary Table 4. DiscoverMP analysis parameters.** The values for threshold 2 (= 0.25) and median filter kernel (= 15) remained constant for all protein samples.
